# Conserved interactions required for in vitro inhibition of the main protease of severe acute respiratory syndrome coronavirus 2 (SARS-CoV-2)

**DOI:** 10.1101/2020.09.10.288720

**Authors:** Alina Shitrit, Daniel Zaidman, Ori Kalid, Itai Bloch, Dvir Doron, Tali Yarnizky, Idit Buch, Idan Segev, Efrat Ben-Zeev, Elad Segev, Oren Kobiler

## Abstract

The COVID-19 pandemic caused by the SARS-CoV-2 requires a fast development of antiviral drugs. SARS-CoV-2 viral main protease (Mpro, also called 3C-like protease, 3CLpro) is a potential target for drug design. Crystal and co-crystal structures of the SARS-CoV-2 Mpro have been solved, enabling the rational design of inhibitory compounds. In this study we analyzed the available SARS-CoV-2 and the highly similar SARS-CoV-1 crystal structures. We identified within the active site of the Mpro, in addition to the inhibitory ligands’ interaction with the catalytic C145, two key H-bond interactions with the conserved H163 and E166 residues. Both H-bond interactions are present in almost all co-crystals and are likely to occur also during the viral polypeptide cleavage process as suggested from docking of the Mpro cleavage recognition sequence. We screened *in silico* a library of 6,900 FDA-approved drugs (ChEMBL) and filtered using these key interactions and selected 29 non-covalent compounds predicted to bind to the protease. Additional screen, using DOCKovalent was carried out on DrugBank library (11,414 experimental and approved drugs) and resulted in 6 covalent compounds. The selected compounds from both screens were tested *in vitro* by a protease activity inhibition assay. Two compounds showed activity at the 50μM concentration range. Our analysis and findings can facilitate and focus the development of highly potent inhibitors against SARS-CoV-2 infection.

## Introduction

The raging pandemic caused by SARS-CoV-2 requires a rapid response of the biomedical community^1,2^. However, novel vaccines and antivirals require time for development, thus repurposing of available drugs is a fast alternative and many attempts using different approaches are made^3–6^. Antiviral drugs are traditionally aimed at viral enzymes and are able to cure or reduce symptoms in several viral infections (HIV, HCV and HSV-1^7^).

SARS-CoV-2, the causative agent of COVID-19, belongs to the genus *Betacoronavirus* and is closely related to SARS-CoV-1, the causative agent of the SARS pandemic outbreak in 2003^8^. Coronaviruses are unsegmented single-stranded positive-stranded RNA viruses, featuring the largest known viral RNA genomes (26 to 32 kilobases in length) infecting humans^9^.

SARS-CoV-2 genome contains 14 open reading frames (ORFs) encoding 27 proteins. First two ORFs at 5’ untranslated region are coding for overlapping polyproteins (replicase 1a (pp1a) and replicase 1ab (pp1ab)) approximately 450kD and 750kD, respectively. The two polyproteins, pp1a and pp1ab, mediate all the functions required for viral replication and transcription. The longer polyprotein (pp1ab) encodes for 15 nonstructural proteins (viral proteins that are not part of the virions) collectively involved in virus replication and possibly in immune evasion.

The functional polypeptides are released from the polyproteins by extensive proteolytic processing. This is primarily achieved by the main protease (Mpro), along with the papain-like protease. Together, they cleave the amino acid backbone at 11 sites on the large polyprotein. This cleavage site involves Leu-Gln↓(Ser/Ala/Gly) sequences (the cleavage site is indicated by ↓)^10^. This cleavage pattern appears to be conserved in the Mpro of SARS-CoV-1.

The Mpro of the coronaviruses is a homodimer. It cleaves the polyprotein using its catalytic dyad that contains the catalytic residues Histidine 41 (H41) and Cysteine 145 (C145) (Fig 1A-C). All of the residues within the active site, including the catalytic residues and adjacent binding residues (polypeptide binding site) belong to one monomer, except for one (Serine 1) from the second monomer^11^.

**Figure 1:**
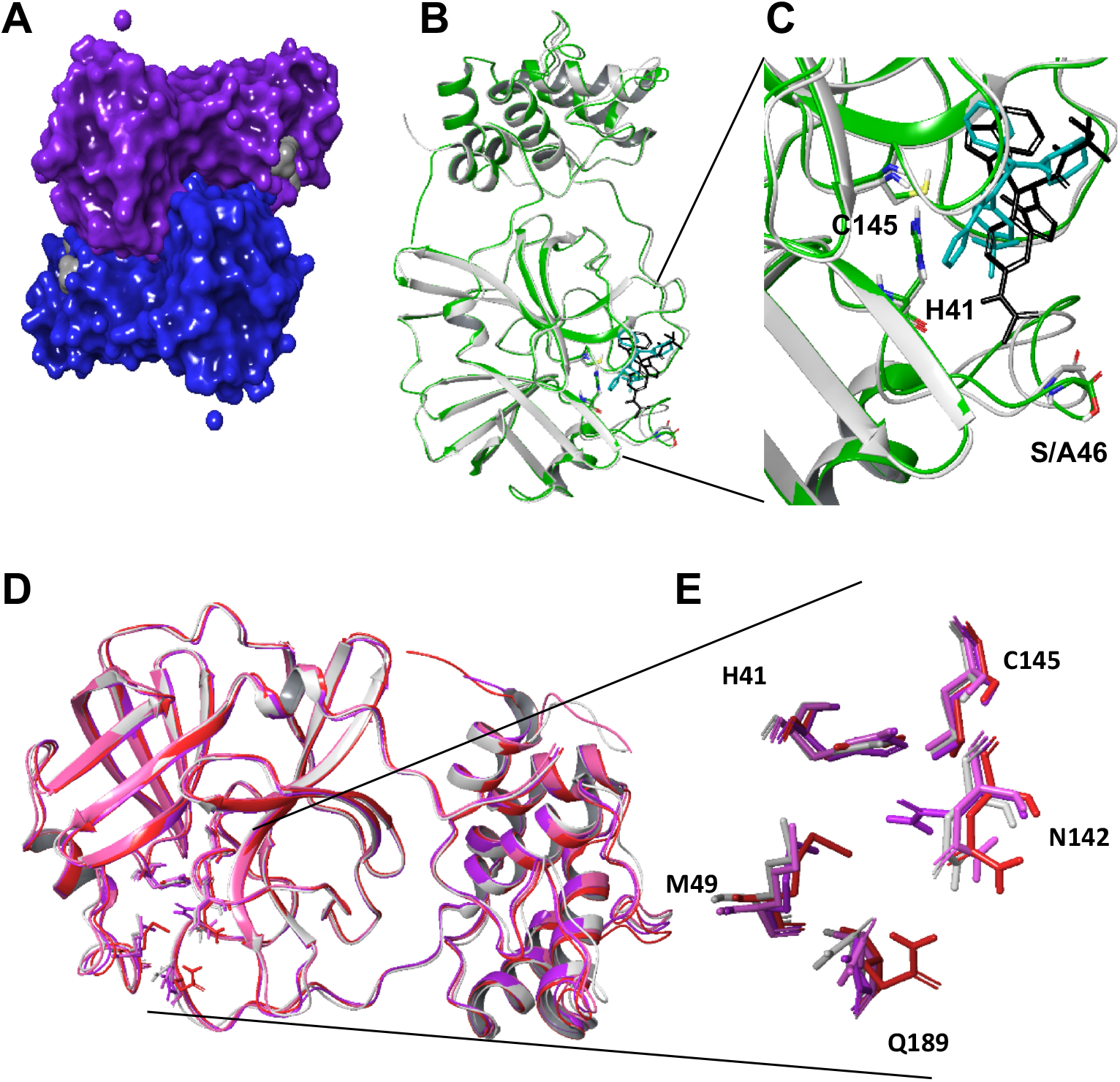
Structural overview of main protease homodimer of SARS-CoV-2 and its binding site. **A.** Surface topology of SARS-CoV-2 Mpro homodimer in complex with the covalent α-ketoamide inhibitor (PDB structure 6Y2F). The two monomers are colored in blue and purple and the inhibitors are represented in gray. **B.** Superimposition of SARS-CoV-2 Mpro (**6W63**, shown as ribbon and colored in green) and SARS-CoV-1 (**4MDS**, shown as ribbon and colored in gray) in complex with their non-covalent inhibitors X77 (shown as sticks and colored in blue) and ML300 (shown as sticks and colored in black), respectively, shown as ribbons. The catalytic residues H41 and C145 are in sticks. The different amino acids SARS-CoV-2 S46 and CoV-1 A46 are shown in sticks. **C.** Magnified view of Figure **1B** (binding site) **D.** Superimposition of the most diverse structures of SARS-CoV-2 and SARS-CoV-1 (available at that time) are shown in ribbons. SARS-CoV-1, **2ZU5** (gray), SFfARS-CoV-2, **5R80** (purple), SARS-CoV-2, **6LU7** (pink), SARS-CoV-2, **6M03** (red), SARS-CoV-2, **6Y2F** (orange). Residues within this site Q189, M49 and N142 and the catalytic residues H41 and C145 are represented in sticks. **E.** Magnified view of Figure **1D**

Several co-crystal structures of the SARS-CoV-2 Mpro were recently solved, enabling the rational design of specific inhibitory compounds^12–15^. The binding site of all the ligands from the co-crystals is found within the Mpro active site. The close relationship of SARS-CoV-2 to SARS-CoV-1 is reflected by high sequence identity of 96.1% and similarity of 99% among their entire proteases protein sequence^16^. In the vicinity of the binding site, the only residue that differs is positioned at residue 46. In SARS-CoV-2 it is a Serine and in SARS-CoV-1 it is an Alanine; however, their side chains point out of the binding site (Figure 1C).

The high similarity between the two viruses’ proteins and the fact that their active sites are practically identical, enable the use of SARS-CoV-1 co-crystals^17–34^ in addition to the available SARS-CoV-2 co-crystals, for understanding the vicinity of the binding site region and defining the important interactions within the SARS-CoV-2 binding site with its inhibitors. In this regard, it was suggested that drugs developed against SARS-CoV-1 might be effective to treat SARS-CoV-2^16^. However, these compounds remained in the preclinical or early clinical stage, without further development into an approved medicine.

In this study, we analyzed the available SARS-CoV-1 and SARS-CoV-2 Mpro co-crystal structures and the developed SARS-CoV-1 inhibitory compounds and identified key interactions required to identify and develop an active inhibitor for the main protease. We conducted a virtual screen using a library of only FDA-approved drugs against SARS-CoV-2 Mpro structure from protein data bank (PDB) [6W63]^13^ using three docking software tools (GOLD^35^ and Glide^36–38^ and DOCKovalent^39^). Several compounds were selected and tested *in vitro* using a protease inhibition assay.

## Results

### Analysis of co-crystals flexibility

To identify the flexibility of the Mpro binding site, we superimposed the SARS-CoV-1 and SARS-CoV-2 apo and co-crystal structures available at the time of our study in the PDB (Table 1). We selected the five most distinct, root-mean-square deviation (RMSD)-wise, structures within the 3D space of the binding site. The selected structures were 2ZU5, 5R80, 6LU7, 6M03, 6Y2F. Three flexible residues within the binding site showed variation in their positions between the different structures: Glutamine 189, Methionine 49 and Asparagine 142 (Figure 1D and E).

**Table 1:**
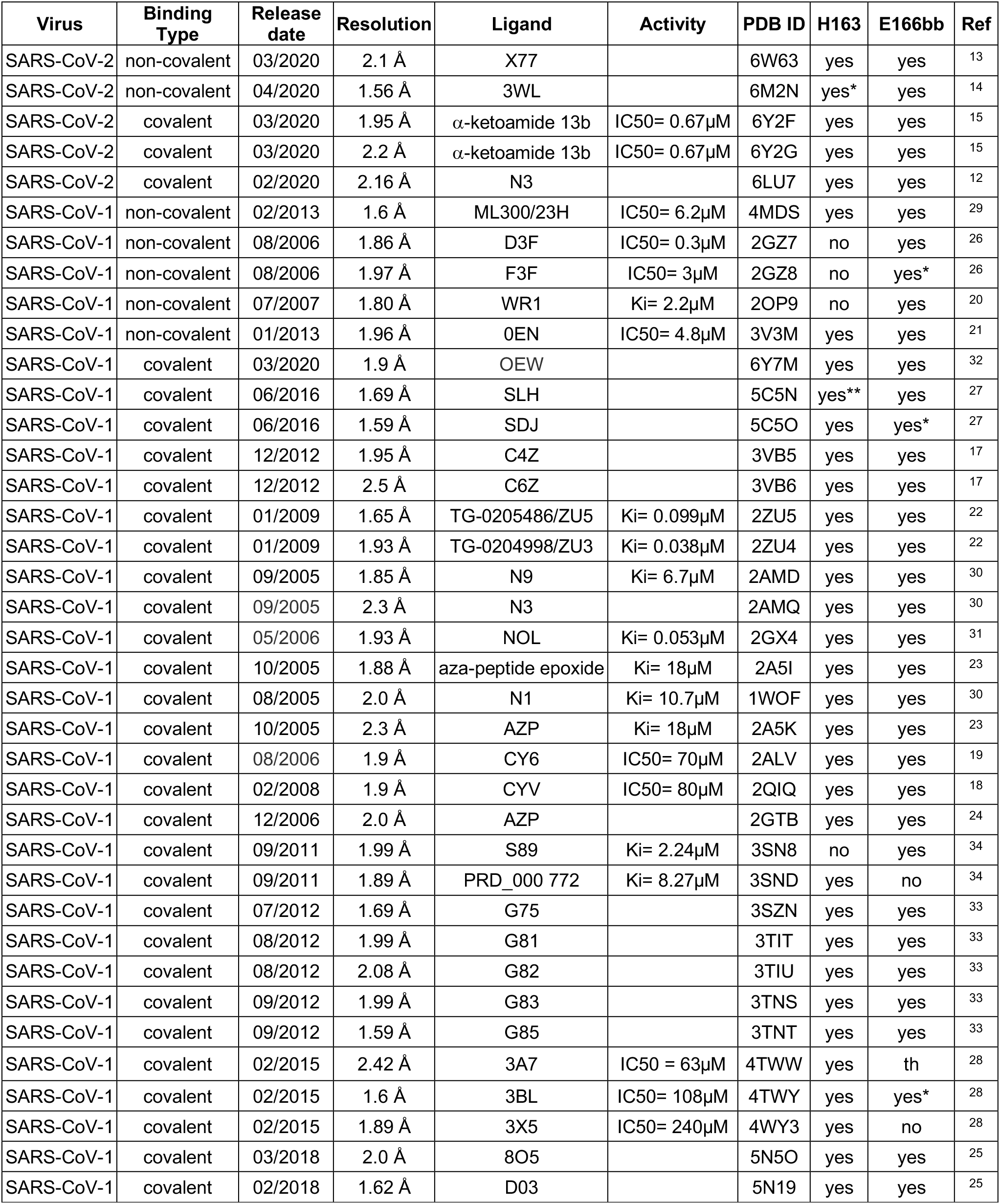
SARS-CoV-1/2 Mpro covalent and non-covalent co-crystals PDB structures. A list of all PDB structures used in this work and their interactions with the two key residues H163 and E166 backbone (bb) are summarized. The known activity from the literature is mentioned when available in either inhibition concentration of 50% (IC50) or inhibitory constant (Ki). *-represent interaction through water molecule **-introduces a donor to the imidazole th-Theoretically represent interaction through water molecule, although the water molecule is not present in the structure.

### Covalent and non-covalent co-crystal interactions

To find the essential interactions required for inhibition of the SARS-CoV-2 Mpro, we analyzed the interactions observed with both covalent and non-covalent inhibitors (Table 1). Most of the co-crystals for SARS-CoV-1 and SARS-CoV-2 contain covalent inhibitors (32 structures). To date, only 6 co-crystals contain non-covalent inhibitors.

Analyzing the co-crystal interactions of the non-covalent inhibitors revealed that the two SARS-CoV-2 co-crystallized ligands, 3WL [6W63] and N3 [6M2N], form H-bonds with protein NH donors: Histidine 163 (H163) and Glutamic acid 166 (E166) backbone. Interestingly, N3 mediates the H-bond interaction with H163 through a water molecule (Figure 2A). In SARS-CoV-1 co-crystals, three out of the four structures showed both H163 and E166 backbone interactions and all exhibited the E166 backbone H-bond interaction. Interestingly, F3F [2GZ8] mediates the E166 backbone interaction through a water molecule.

**Figure 2:**
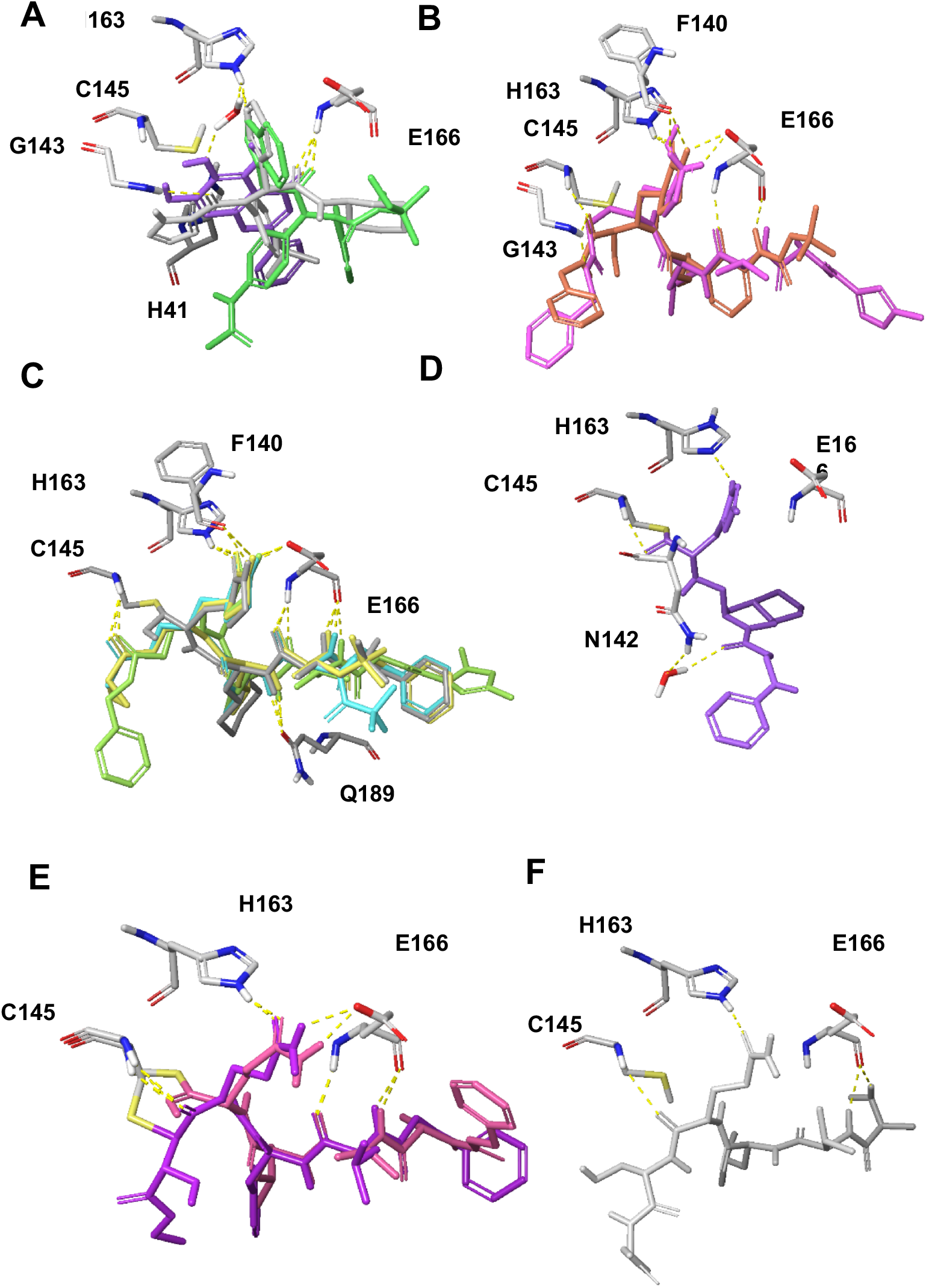
Co-crystals interactions and important residues. **A.** Superimposition of non-covalent co-crystals 6W63 (gray), 4MDS (green) and 6M2N (purple) with protein residues shown in sticks (colored by element). **B.** Superimposition of covalent co-crystals 6LU7 (pink), 6Y2F (orange). **C.** Superimposition of covalent co-crystals 2GX4 (gray), 2ZU4 (cyan), 2AMQ (green), 2ZU5 (yellow), shown in sticks. **D.** 5C5N (purple), shown in sticks. Protein interactions residues are in shown in sticks. **E.** Two co-crystals with peptides 2A5I (purple), 3VB5 (pink). **F.** The recognition sequence peptide docked within the 6LU7 (gray).

Additional interactions were observed with the following amino acids: The catalytic H41 with D3F [2GZ7] and ML300 [4MDS]. The catalytic C145 forms a H-bond with F3F [2GZ8]. G143 backbone with X77 [6W63] and N142 and F140 with F3F [2GZ8]. Many hydrophilic moieties of the ligands are surrounded by water molecules that mediate the interaction of the inhibitor with the protein.

All covalent compounds interact with the catalytic C145 in the co-crystals. Interestingly, most (31 out of 32) comprise also a non-covalent interaction, H-bond with H163 similarly to the non-covalent compounds. All of the covalent and non-covalent inhibitors present a H-bond acceptor to the side chain imidazole ring of H163 (see for example Figure 2A-C). The only exception, presenting a H-bond donor towards H163, is SLH inhibitor [5C5N] (Figure 2D). Since the hydrogen can reside on either nitrogen, (N1-H or N3-H tautomers) it interacts with the imidazole N acceptor. E166 backbone that is interacting with all non-covalent ligands is also a key residue for most covalent ligands. Most ligands (30 out of 32) form H-bond interactions with the E166 backbone NH and some form an additional interaction with the E166 backbone carbonyl oxygen (for example, α-ketoamide 13b [6Y2F], ZU3 [2ZU4], N3 [2AMQ and 6LU7] and ZU5 [2ZU5], Figure 2B and C). Few structures mediate E166 backbone interaction through a water molecule (for example, F3F [4TWY] and SDJ [5C5O]) (Table 1). In addition to E166 backbone interactions, some ligands interact with E166 side chain either via salt bridge (for example, N3 [6LU7] and SLH [5C5N]) or through a H-bond interaction (for example, α-ketoamide 13b [6Y2F] and ZU3 [2ZU4], Fig 2B and 3C).

**Figure 3:**
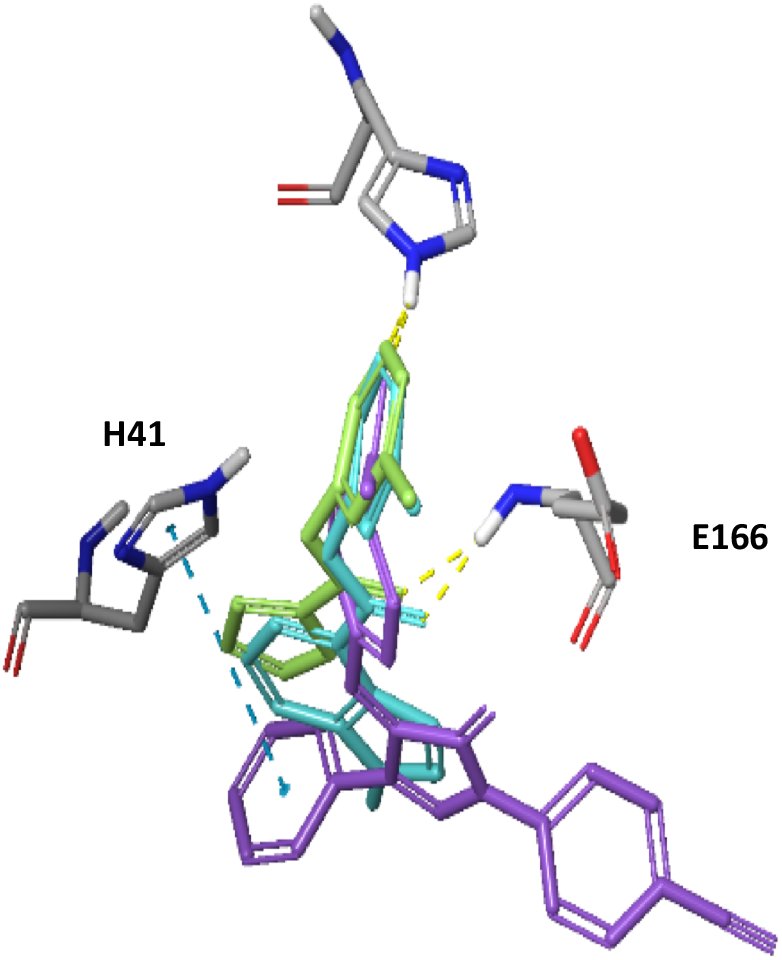
SARS-CoV-1 developed inhibitor docked to Mpro 6W63 PDB structure. Zhang2007 cmp37 (green), ghosh2008 cmp10 (cyan), Lu2006_Pyrazolone cmpd2p (purple). Important residues for interactions are shown in sticks.

Other backbone interactions can be detected with G143 [6LU7, 6Y2F and 2ZU4], H164 [6LU7] and F140 [2GX4 and 6LU7]. In addition, side chain interaction is formed by ZU3 [2ZU4] with the flexible residue Q189 (Figures 2B and C).

### Docking of the cleavage recognition sequence

The proteolytic activity of Mpro catalyzes cleavage between Serine and Glutamine within the viral polypeptides. To characterize the interactions required for cleavage, we analyzed two co-crystals with peptidomimetic inhibitors [2A5I, 3VB5]. In these structures, the side chain of the catalytic C145 binds the peptide Serine backbone. The catalytic C145 side chain is rigid in all co-crystals except for [2A5I] in which the side chain adopts a unique conformation. Interestingly, the peptide Glutamine side chain of the cleavage site is anchored by a H-bond interaction with the H163 imidazole (Fig 2E).

To characterize the interactions of the cleavage recognition sequence peptide we chose, based on the peptidomimetic inhibitors, the following sequence: Ala, Val, Leu, Gln, Ser, Ala, Gly. We docked (using Glide) the recognition sequence peptide to SARS CoV-2 6LU7 crystal structure and superimposed the two co-crystals with the peptides [2A5I, 3VB5]. The Glutamine within the recognition sequence adopted the same conformation as in the two co-crystals, presenting the same H-bond interaction with H163 imidazole (Fig 2F). In addition, the peptide’s Valine and Alanine backbone interact with the E166 backbone (through water molecules).

### Docking of known SARS-CoV-1 Mpro inhibitors

Since the SARS-CoV-1 outbreak in 2003, several studies have developed inhibitors for Mpro of SARS-Cov-1^40^. To verify that our observed interactions are required for Mpro inhibition, we docked non-covalent SARS-CoV-1 Mpro inhibitors to the SARS-CoV-2 Mpro binding site [6W63]. The same two common interactions (H163 and E166) were present in all compounds tested (see for example few known inhibitors in Figure 3).

In summary, the two hydrophilic interactions with H163 and E166 backbone exist in most of the covalent and non-covalent co-crystal ligands and all of these co-crystals show at least one of these interactions. The known inhibitors show the same pattern of interactions and these interactions seem to play a role in the recognition sequence binding, thus highlighting them as biologically significant. Therefore, in the screening process these interactions were chosen as filtering criteria, allowing to pass only poses that satisfied at least one of these two interactions, for further analysis.

In addition, the Schrödinger SiteMap tool^41,42^ identified two hydrophobic regions within the vicinity of the binding site and we found that most of the covalent and non-covalent co-crystal ligands and the known inhibitors introduced hydrophobic moieties within those regions.

### Non-covalent docking using GOLD and Glide

To identify possible inhibitors from the FDA-approved drugs we used [6W63] protein structure as a template for virtual screening, applying two docking software (as recommended^43^). The prepared ligand set originating from the ChEMBL drug database was docked either using GOLD, outputting 10 conformations (poses) for each compound resulting in 46,190 poses (3,634 unique drugs), or using Glide, outputting at most 5 conformations (poses) per compound resulting in 22,004 poses (3,620 unique drugs).

We filtered the poses based on the two significant interactions identified in our analysis: H163 imidazole H-bond and E166 backbone amine H-bond (see the Materials and Methods section for details). We chose the best docking poses that satisfied either one or both of these interactions, resulting in at most three poses for each compound. This stage resulted in 2,993 unique compounds poses in GOLD and 1,969 unique compounds in Glide. We manually selected the filtered poses resulting in 21 compounds in GOLD and 13 in Glide. Altogether, a total of 29 unique compounds (4 of which were selected in both methods) were selected and sent for assessment using the protease inhibition assay. One compound, selected by the GOLD software, GSK-256066, showed 37% inhibition at concentration of 50μM (Supplementary Table 1).

### Covalent docking using DOCKovalent

Several covalent docking software were developed at Nir London’s lab at the Weizmann institute ^39^. As there are very few possible known drugs that can perform covalent binding, we used preclinical and clinical compounds from the DrugBank database ^44^. This database was filtered to contain only compounds with covalent warheads that can be docked using DOCKovalent (see Materials and Methods) to: [6M03, 5R7Y, 5R7Z, 6Y2F, 6W63, 4MDS, 2GX4, 6LU7] PDB structures. These compounds were visually inspected and we selected the ones that showed additional interactions to the C145 covalent interaction. We tested 5 nitriles and one Michael acceptor and two of the nitriles (bicalutamide and ruxolitinib) showed 36% and 20% inhibition at 50μM, respectively (Supplementary Table 2)

## Discussion

Antiviral drugs targeting the Mpro of SARS-CoV-2 could support the fight against the global COVID-19 pandemic. Here, to identify possible inhibitors of the SARS-CoV-2 Mpro, we have explored the co-crystal structures of the Mpro proteins of SARS-CoV-2 and SARS-CoV-1. We identified two common interactions involving H163 and E166 that appeared in most co-crystals. We screened *in silico* drug databases for covalent and non-covalent compounds. Possible compounds were further tested in a protease inhibition assay and we found several compounds that reduce protease activity by more than 30%.

The Mpro protein sequence of SARS-CoV-2 is highly similar (99%) to SARS-CoV-1. In the region of the binding site only one residue is different. Some studies suggested that the differences between the two proteins affected the ability to bind inhibitors^45,46^. On the other hand, several studies and our protease inhibition assay show that inhibitors identified for SARS-CoV-1 Mpro also inhibit SARS-CoV-2 Mpro (see Supplementary Table 3). Further co-crystals of SARS-CoV-1 [2MAQ] and SARS-CoV-2 [6LU7] Mpro with the identical inhibitor (N3) show similar interactions with the protease binding site^12^. Thus, we inferred that the binding to the binding site of both viruses is comparable and therefore we were able to analyze the key interactions based on co-crystals obtained from both viruses.

We identified that all co-crystals have at least one of two key interactions with H163 and E166. Docking of the recognition sequence peptide into the binding site revealed that H163 and E166 form H-bonds with the peptide. Specifically, the imidazole ring of H163 interacts with the conserved Glutamine of the cleavage site^11^ while E166 interacts with the Alanine and Valine from the recognition sequence. Interestingly, E166 side chain interacts with Serine 1 NH_2_-terminal of the second monomer^11,47^. This salt bridge interaction minimizes the conformational flexibility of E166 backbone and assists in generating the correct orientation of the substrate binding site, which explains the importance of dimerization for the catalytic activity^47^. H163 and E166 amino acids are conserved among all human coronaviruses (2 alpha- and 5 beta-coronaviruses Figure 4), unlike H164 and Q189 that were previously identified as important interactions of several inhibitors^12,22^. Thus, drugs developed to interact with these amino acids may be effective against all human coronaviruses and could potentially prevent the emergence of viral resistance.

**Figure 4:**
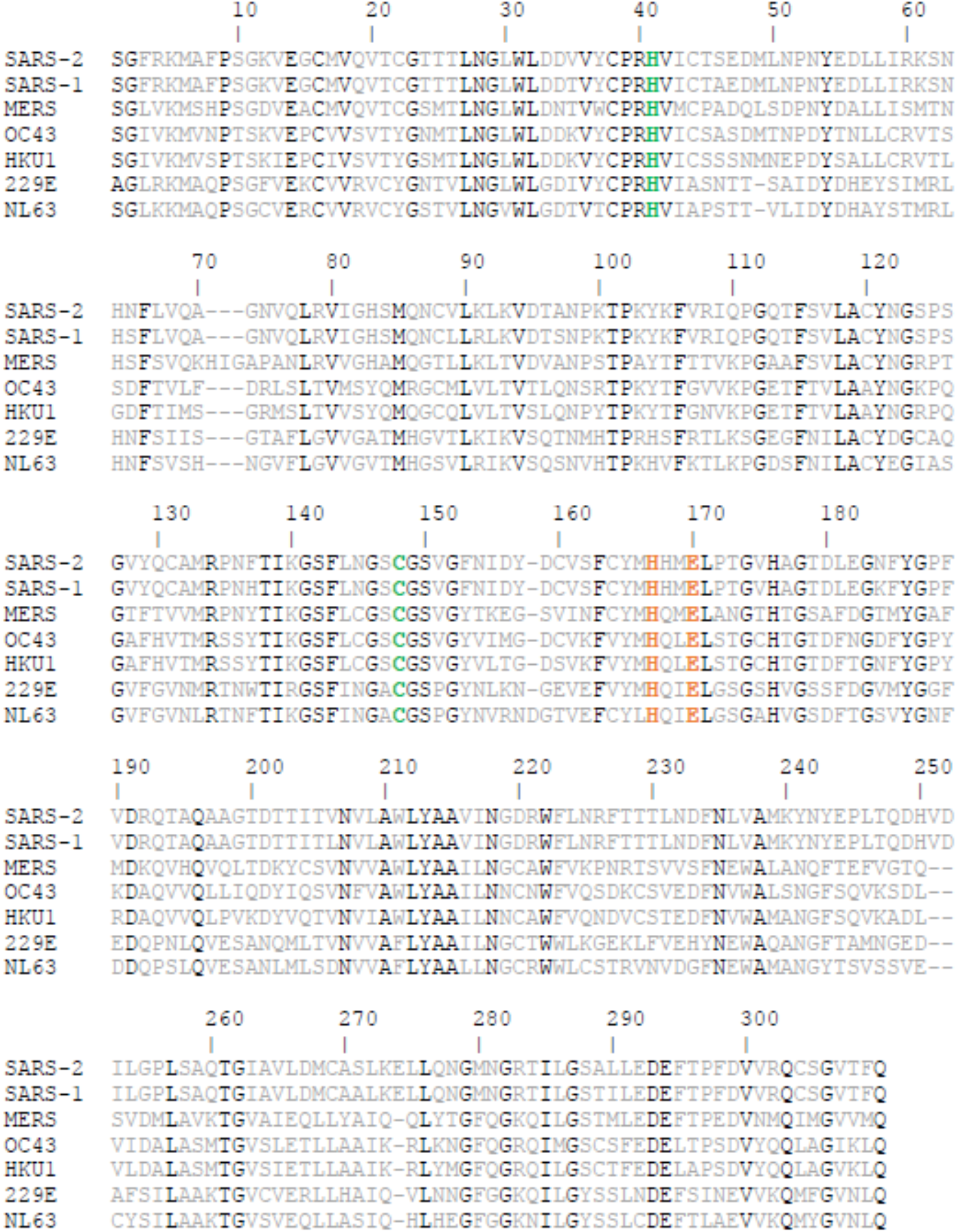
Conservation of human coronaviruses Mpro. Conserved residues are colored in black. Specifically, catalytic dyad residues (H41 and C145) are colored in green. H163 and E166 are colored in orange.

Several attempts to identify in-silico inhibitors of SARS-CoV-2 Mpro have been already published^48–52^. All of these did not validate their virtual screen results by in-vitro experiments. Further, these studies used either [6LU7] or [6Y2F], as their template for the computational screening. We used [6W63] as the protein structure for our non-covalent docking, as [6W63] ligand is non-covalent while [6LU7] and [6Y2F] ligands are covalent. The protein structure of [6W63] differs from [6LU7] and [6Y2F] in the identified co-crystals flexibility residues M49, Q189 and N142 (Figure 1D and E). For the covalent docking we used seven different crystal structures (see results) to allow more flexibility in the binding site.

Our two screening analyses resulted in two clinically approved drugs that inhibit the Mpro by over 30% in 50μM.: The first one is GSK-256066, a phosphodiesterase (PDE) 4 inhibitor^53^ that was under development in phase 2 for the treatment of chronic obstructive pulmonary disease (COPD), asthma and seasonal allergic rhinitis. It is administered as an inhalation formulation (powder) and as an intranasal formulation (nasal spray suspension). Our model suggests that GSK-256066 forms a H-bond with H163 and additional two H-bonds with the amine and carbonyl of E166 backbone (Figure 5). It inhibits the Mpro by 37% at a concentration of 50μM.

**Figure 5.**
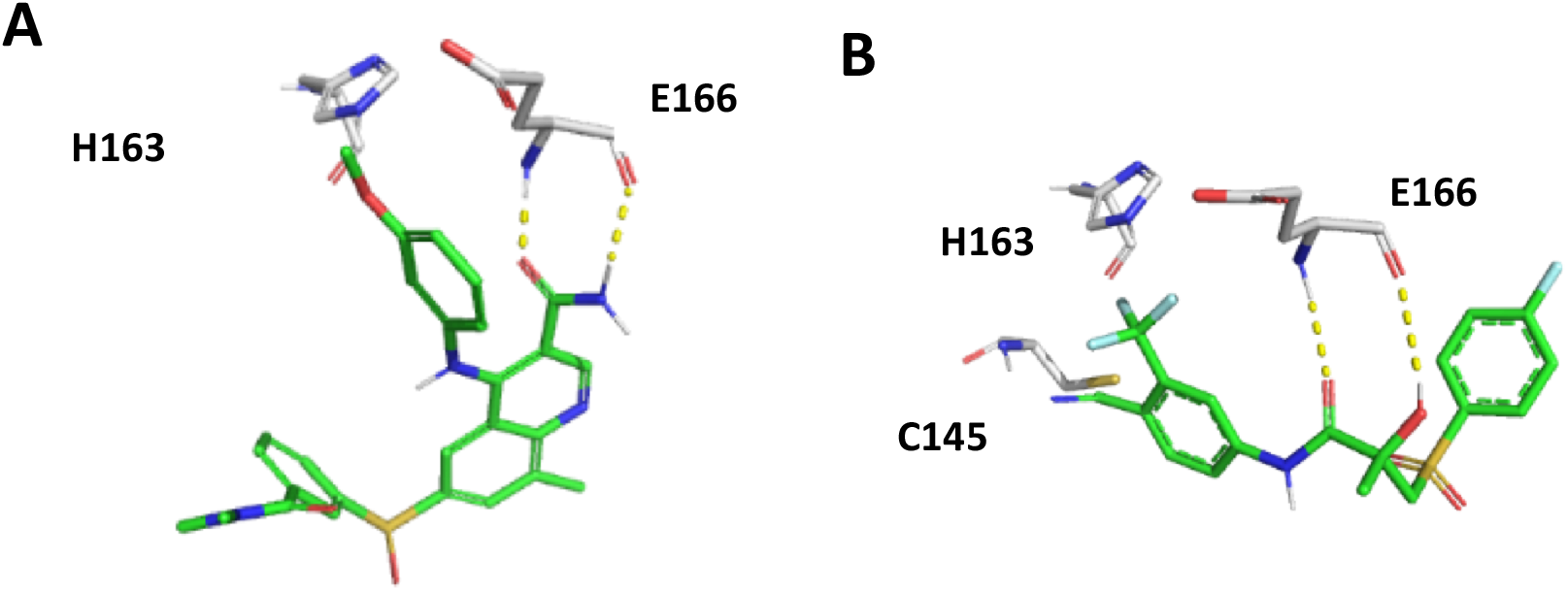
FDA-approved drugs that inhibit Mpro. A. GSK-256066 (colored by element) B. Bicalutamide (colored by element). Important residues for interactions are shown in gray element sticks.

Another drug that showed inhibition of the Mpro is bicalutamide, which was selected from the covalent screening. It contains an aryl nitrile that can covalently bind to the protein. Bicalutamide is an oral non-steroidal anti-androgen for prostate cancer. It is comprised of a 50:50 racemic mixture of the (*R*)- and (*S*)-enantiomers. Bicalutamide binds to the androgen receptor. Our model suggests that its nitrile group covalently binds to C145 and forms two H-bonds with the amine and carbonyl of the E166 backbone (Figure 5). Bicalutamide was tested in two experiments and inhibited Mpro by 37% and 33% at a concentration of 50μM.

Several compounds that were previously identified as inhibitors with sub-micromolar potency were active in our protease inhibition assay (Supplementary Table 3). Two of these inhibitors with known sub-micromolar activity, showed limited inhibition (39% and 9%) at a concentration of 50μM in our protease activity assay (Supplementary Table 3). Thus, GSK-256066 and bicalutamide, that were identified in our protease inhibition assay, have a similar inhibitory activity at the same concentration. These results suggest that more assays should be conducted to test repurposing of these drugs as anti-SARS therapeutics.

In conclusion, our analysis of the structural constraints required for the inhibition of SARS-CoV-2 Mpro has suggested key interactions with several amino acids in the active pocket of the protein. We were able to identify several approved drugs with a potential to inhibit Mpro activity, indicating that our analysis could be used for virtual screenings and rational drug development.

## Materials and Methods

### Protein data bank (PDB) search

The protein data bank was searched for SARS-CoV/SARS-CoV-1/SARS-CoV-2 Mpro. Non SARS-CoV structures and non-human SARS-CoV like structures were omitted. Co-crystals binding fragments were not added to this analysis due to their non-drug like structures. We anticipate that few of the available structures might be overlooked using these search criteria. All PDB structures found and analyzed are mentioned in Table 1. Throughout the text, PDB IDs are marked with square brackets.

### Preparing a drug library from ChEMBL for non-covalent docking

The ChEMBL database contains 6,900 drugs in various stages of clinical trials. To focus our computational screen, the following filters were applied: small molecules that were at clinical phase 2 or higher, number of rotatable bonds < 14, MW freebase 200-990, ATC Class Level 1: all except D-Dermatologics and V-Various. This filtering resulted in a 4,239-compound library. To prepare the ligands for docking simulations, LigPrep (Schrödinger Release 2020-1: LigPrep, Schrödinger, LLC, New York, NY, 2020) was applied on the exported library. The following settings were used: (a) The OPLS3e force field was chosen^54^; (b) Possible protonation states were generated using Ionizer at a target pH of 7.4; (c) All ligands were desalted; (d) No tautomers were generated; (e) At most two stereoisomeric forms were produced per ligand for unspecified chiral centers. These constraints enabled us to expand the initial 4,239-compound library to only 4,623 ligands.

### Preparing a drug library from DrugBank for covalent docking

We used the DrugBank database^44^ that includes 11414 preclinical and clinical small molecules. These compounds were filtered by ≤500D MW and ≤5 rotatable bonds. Only compounds that contain covalent warheads (Michael acceptors: O=CC=[C;H1,H2] or nitriles) were selected as they can covalently bind the thiol of C145. The filtering resulted in a library of 437 ligands (258 Michael acceptors and 179 nitriles).

### Docking

The 4,623 prepared ligands originating from ChEMBL were docked using Glide (Schrödinger Release 2020-1: Glide, Schrödinger, LLC, New York, NY, 2020) to the Mpro structure from [6W63], keeping the protein structure rigid and ligands flexible, with no constraints applied on specific receptor-ligand hydrogen bond (H-bond) interactions. The standard precision (SP) mode of Glide was used based on the OPLS3e force field, writing out at most 5 poses per ligand. The 22,003 conformations (poses) were filtered by requiring at least one of the two key H-bond interactions with H163 and E166. The default maximum H-bond distance criterion of 2.5 Å was stretched to 3.0 Å. This filter resulted in three groups of poses: (a) 517 poses interacting with both H163 and E166; (b) 2,088 poses forming a H-bond with H163 only; and (c) 2,678 poses forming a H-bond with E166 only. In each of the filtered groups, the pose with the best Glide score per each ligand was selected, resulting in 293, 879 and 1,347 poses, respectively. We further narrowed down the number of poses by eliminating drugs with molecular charge below −1 using Maestro’s Ligand Filtering utility. Applying this filter resulted in 260, 820 and 1,283 poses, respectively. Removing the duplicate poses (i.e. those overlapping with one associated with either or both of the two other groups) resulted in a total of 1,969 unique poses.

The 4,239 ChEMBL-derived (“pre-Ligprep”) drugs were also docked using GOLD Standard docking^35^ to Mpro structure from [6W63] resulting in 46,190 poses (10 poses per ligand). Identical filters as in Glide docking were applied resulting in a total of 2,993 unique poses that were grouped by interactions to (a) 86 poses interacting with both H163 and E166; (b) 1,011 poses forming H-bond with H163 only; and (c) 1,896 poses forming H-bond with E166 only.

### Selection

In our manual selection we preferred ligands that in addition to one or two important interactions (H163 and E166) also formed interactions with additional residues that were found in the co-crystal structure (for example Gly143 backbone). In addition, we favored compounds that did not violate the two hydrophobic regions within the binding site as calculated by Maestro’s SiteMap tool (Schrödinger Release 2020-1: SiteMap, Schrödinger, LLC, New York, NY, 2020.^41,42^).

### Protease inhibitor activity assay

35 compounds were obtained as detailed in Supplementary Table 1 and 2. The compounds were prepared in assay ready plates (Greiner 784900) using Labcyte Echo 555 and diluted in DMSO to concentration of 0.5%. 5nM Mpro and 375nM [5-FAM]-AVLQSGFR-[Lys(Dabcyl)-K-amide substrate (in 20mM HEPES pH=7.3, 50mM NaCl, 10% Glycerol, 0.01% Tween-20, 1mM TCEP) were added to the compounds and incubated for 30 minutes at room temperature. Fluorescence was read at 480/520 ex/em in BMG Pherastar FS.

The Mpro inhibition assay was carried out in the Mantoux Bioinformatics institute of the Nancy and Stephen Grand Israel National Center for Personalized Medicine (INCPM), Weizmann Institute of Science.

## Funding

This work was supported by grants from the Israel Science foundation (grant #1387/14) to O.K. The funders had no role in study design, data collection and analysis, decision to publish, or preparation of the manuscript.

## Author Contributions

AS and OKo designed research; AS, DZ, OKa, IB, DD, TY, IB, IS, EBZ and ES analyzed data; AS, DZ, OKa, IB, DD, TY, IB, IS, EBZ and ES performed research; AS and OKo wrote the paper.

## Acknowledgement

We thank Craig Coel and Katalin Phimister from Schrödinger for all their support, Haim Barr and Nir London for their help and all Kobiler lab members for their comments.

## Supplementary Table Legends

**Supplementary Table 1: Non-covalent compounds that were selected for testing and their % of inhibition at 50μM concentration.** A list of all non-covalent compounds tested in the protease inhibition assay after selection either by the GOLD, Glide or both docking tools. Percent average inhibition at 50μM is presented (Avg. Inh).

**Supplementary Table 2: Covalent compounds that were selected for testing and their % of inhibition at 50μM concentration.** A list of all covalent compounds tested in the protease inhibition assay after selection. Percent average inhibition at 50μM is presented (Avg. Inh).

**Supplementary Table 3: Several SARS-CoV-1 known Mpro inhibitors and their results in the protease inhibition assay.** The structures (column A) and activity Column C) of several SARS-CoV-1 inhibitors are known from the publications (column D) along with the results obtained in our protease inhibition assay (columns E-H).

## Notes

### Competing Interest Statement

The authors have declared no competing interest.

