## Supplemental tables for "Conserved interactions required for in vitro inhibition of the main protease of severe acute respiratory syndrome coronavirus 2 (SARS-CoV-2)"

### Supplementary Table 1

| Drug name | Structure | Smiles | Glide/ GOLD | Avg Inh. (%) |
|---|---|---|---|---|
| **GSK-256066** | 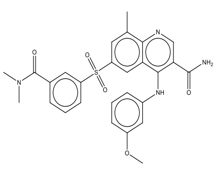 | CN(C)C(=O)c1cc(ccc1)S(=O)(=O)c(c2)cc(C)c(c23)ncc(C(=O)N)c3Nc4cc(OC)ccc4 | GOLD | 36.96 |
| **AVANAFIL** | 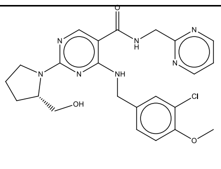 | n1cccnc1CNC(=O)c2c(NCc3cc(Cl)c(cc3)OC)nc(nc2)N4CCC[C@H]4CO | GOLD | 12.24 |
| **LAMIVUDINE** | 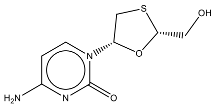 | OC[C@@H]1O[C@@H](CS1)n(c(n2)=O)ccc2N | Glide | 9.29 |
| **BIMIRALISIB** | 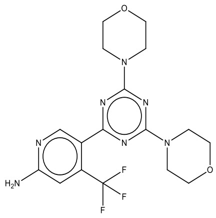 | c1c(N)ncc(c1C(F)(F)F)-c2nc(N3CCOCC3)nc(n2)N4CCOCC4 | GOLD | 5.97 |
| **MELOXICAM** | 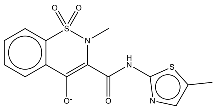 | Cc1cnc(s1)NC(=O)C(=C2[O-])N(C)S(=O)(=O)c(c23)cccc3 | GOLD | 4.82 |
| **VOXTALISIB** | 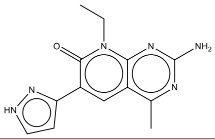 | Cc1nc(N)nc(c12)n(CC)c(=O)c(c2)-c3cc[nH]n3 | GOLD | 4.38 |
| **BENZNIDAZOLE** | 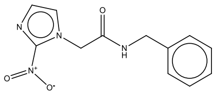 | [O-][N+](=O)c1nccn1CC(=O)NCc2ccccc2 | GOLD | 4.09 |
| **TAK-715** | 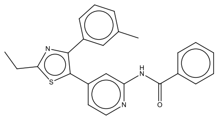 | c1ccccc1C(=O)Nc(ncc2)cc2-c(sc(n3)CC)c3-c(cc4C)ccc4 | Both | 2.47 |
| **DARUSENTAN** | 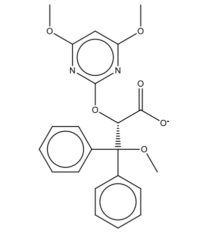 | c1ccccc1C(OC)(c2ccccc2)[C@@H](C([O-])=O)Oc(n3)nc(OC)cc3OC | Both | 2.42 |
| **ERLOTINIB** | 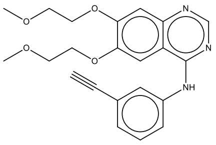 | C#Cc1cc(ccc1)Nc2ncnc(c23)cc(OCCOC)c(c3)OCCOC | Glide | 2.10 |

| | | | | |
|---|---|---|---|---|
| **TAMOXIFEN CITRATE** | 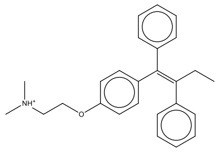 | c1ccccc1C(\CC)=C(c2ccccc2)/c3ccc(cc3)OCC[NH+](C)C | GOLD | 1.61 |
| **SOLCITINIB** | 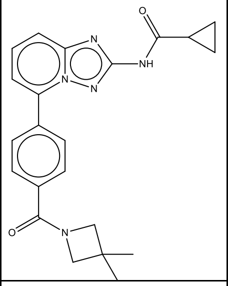 | C1CC1C(=O)Nc(n2)nn(c23)c(ccc3)-c(cc4)ccc4C(=O) N(C5)CC5(C)C | GOLD | 0.84 |
| **PAGOCLONE** | 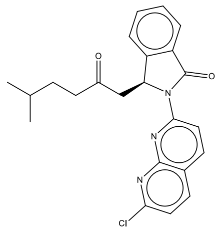 | CC(C)CCC(=O)C[C@@H](c(c12)cccc2)N(C1=O)c(cc3)nc(c34)nc(Cl)cc4 | GOLD | <0 |
| **VIPADENANT** | 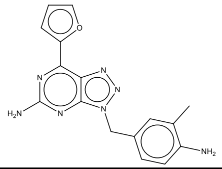 | c1cc(N)c(C)cc1Cn(nn2)c(c23)nc(N)nc3-c4ccco4 | GOLD | <0 |
| **VADADUSTAT** | 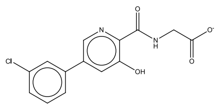 | [O-]C(=O)CNC(=O)c1c(O)cc(cn1)-c2cc(Cl)ccc2 | GOLD | <0 |
| **CEPHALEXIN** | 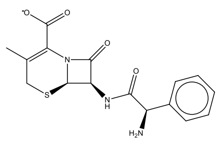 | c1ccccc1[C@@H](N)C(=O)N[C@@H]2C(=O)N([C@@H]23)C(C([O-])=O)=C(C)CS3 | Glide | <0 |
| **ARMODAFINIL (MBX-102)** | 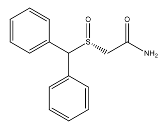 | C1=CC=C(C=C1)C(C2=CC=CC=C2)[S@](=O)CC(=O)N | Glide | <0 |
| **TERIFLUNOMIDE** | 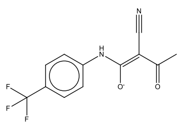 | CC(=O)C(\C#N)=C([O-])\Nc(cc1)ccc1C(F)(F)F | Glide | <0 |
| **BETRIXABAN** | 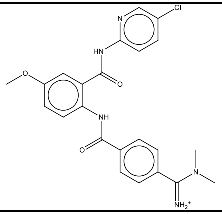 | CN(C)C(=[NH2+])c1ccc(cc1)C(=O)Nc(ccc(c2)OC)c2C(=O)Nc(nc3)ccc3Cl | GOLD | <0 |
| **RIBOCICLIB** | 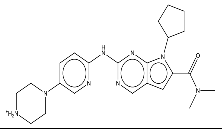 | C1CCCC1n(c(c2)C(=O)N(C)C)c(c23)nc(nc3)Nc(nc4)ccc4N5CC[NH2+]CC5 | GOLD | <0 |

| | | | | |
|---|---|---|---|---|
| **SELICICLIB** | 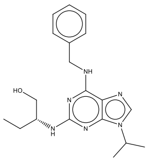 | CC(C)n(cn1)c(c12)nc(N[C@H](CC)CO)nc2NCc3ccccc3 | Both | <0 |
| **LAROTRECTINIB** | 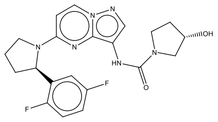 | O[C@H]1CCN(C1)C(=O)Nc2cnn(c23)ccc(n3)N4CCC[C@@H]4c5c(F)ccc(F)c5 | Glide | <0 |
| **GLAFENINE** | 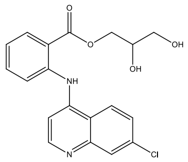 | C1=CC=C(C(=C1)C(=O)OCC(CO)O)NC2=C3C=CC(=CC3=NC=C2)Cl | Both | <0 |
| **TG100-115** | 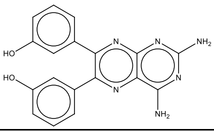 | Nc1nc(N)nc(c12)nc(-c3cc(O)ccc3)c(n2)-c4cc(O)ccc4 | GOLD | <0 |
| **DAROLUTAMIDE** | 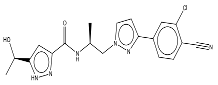 | N#Cc1c(Cl)cc(cc1)-c2ccn(n2)C[C@H](C)NC(=O)c3cc([nH]n3)[C@@H](C)O | Glide | <0 |
| **REBAMIPIDE** | 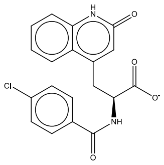 | c1cc(Cl)ccc1C(=O)N[C@H](C([O-])=O)Cc2cc(=O)[nH]c(c23)cccc3 | GOLD | <0 |
| **LONIDAMINE** | 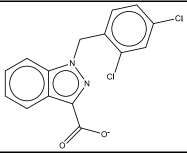 | c1cccc(c12)n(nc2C([O-])=O)Cc3c(Cl)cc(Cl)cc3 | Glide | <0 |
| **GANDOTINIB** | 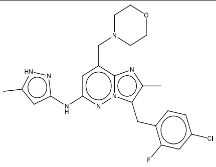 | c1cc(Cl)cc(F)c1Cc(c(n2)C)n(c23)nc(Nc(n[nH]4)cc4C)cc3CN5CCOCC5 | GOLD | <0 |
| **RIBOFLAVIN** | 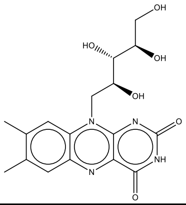 | O=c1[nH]c(=O)nc(c12)n(C[C@H](O)[C@H](O)[C@H](O)CO)c3c(n2)cc(C)c(C)c3 | Glide | <0 |

### Supplementary Table 2

| Drug name | Structure | Smiles | Covalent moiety | Avg Inh. |
|---|---|---|---|---|
| **Bicalutamide** | 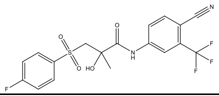 | CC(O)(CS(=O)(=O)C1=CC=C(F)C=C1)C(=O)NC1=CC(=C(C=C1)C#N)C(F)(F)F | Nitril | 35.01 |
| **Ruxolitinib** | 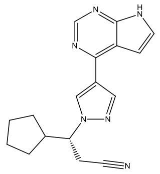 | N#CC[C@H](C1CCCC1)N1C=C(C=N1)C1=C2C=CNC2=NC=N1 | Nitril | 19.23 |
| **Trilostane** | 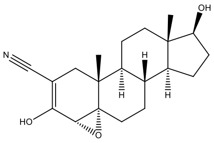 | [H][C@@]12CC[C@H](O)[C@@]1(C)CC[C@@]1([H])[C@@]2([H])CC[C@@]23O[C@@H]2C(O)=C(C[C@]13C)C#N | Nitril | 0.37 |
| **Anastrozole** | 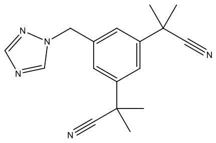 | CC(C)(C#N)C1=CC(=CC(CN2C=NC=N2)=C1)C(C)(C)C#N | Nitril | 4.07 |
| **WP1066** | 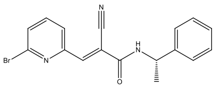 | C[C@H](NC(=O)C(=C\C1=CC=CC(Br)=N1)\C#N)C1=CC=CC=C1 | Michael Acceptors | <0 |
| **Tofacitinib** | 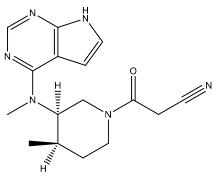 | [H][C@@]1(C)CCN(C[C@]1([H])N(C)C1=NC=NC2=C1C=CN2)C(=O)CC#N | Nitril | <0 |

### Supplementary Table 3

| Structure | Smiles | Name as in reference | IC50 (μM) as in reference | Avg Inh. at 20μM | Avg Inh. at 50μM | IC50 (μM) | <u>Relative Solubility at 20mM</u> |
|---|---|---|---|---|---|---|---|
| 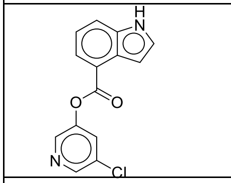 | O=C(c(ccc1)c2c1[nH]cc2)Oc3cc(Cl)cnc3 | 10_GRL-0496[1] | 0.03 | 99.84 | 100.01 | 0.05 | 0.99 |
| 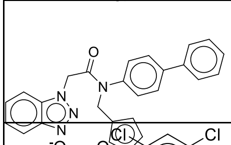 | O=C(N(c1ccc(c2ccccc2)cc1)Cc3cscc3)Cn(nn4)c5c4cccc5 | 17a[1] | 0.051 | 30.41 | 39.14 | 5.32 | 0.64 |
| 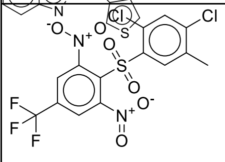 | Cc1c(Cl)cc(Cl)c(S(=O)(c2c([N+]([O-])=O)cc(C(F)(F)F)cc2[N+]([O-])=O)=O)c1 | 3[2] | 0.3 | | 9.61 | | |
| 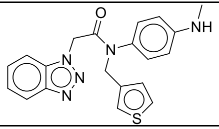 | CNc1ccc(N(C(Cn(nn2)c3c2cccc3)=O)Cc4cscc4)cc1 | 16j[3] | 2.1 | 57.12 | 82.62 | 4.33 | 0.99 |
| 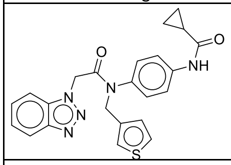 | O=C(C1CC1)Nc2ccc(N(C(Cn(nn3)c4c3cccc4)=O)Cc5cscc5)cc2 | 16e (ML300)[3] | 4.11 | 68.97 | 70.20 | 3.99 | 0.90 |
| 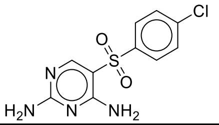 | Nc1nc(N)c(S(=O)(c2ccc(Cl)cc2)=O)cn1 | 5[2] | 6 | | -22.07 | | |
| 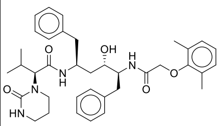 | Cc1c(OCC(N[C@H]([C@H](C[C@@H](NC([C@@H](N2CCCNC2=O)C(C)C)=O)Cc3ccccc3)O)Cc4ccccc4)=O)c(C)ccc1 | Lopinavir[4] | >32 | 7.22 | 12.97 | | 0.83 |


1 Ghosh, A. K. *et al.* Design, synthesis and antiviral efficacy of a series of potent chloropyridyl ester-derived SARS-CoV 3CLpro inhibitors. *Bioorg Med Chem Lett* **18**, 5684-5688, doi:10.1016/j.bmcl.2008.08.082 (2008).

2 Lu, I. L. *et al.* Structure-based drug design and structural biology study of novel nonpeptide inhibitors of severe acute respiratory syndrome coronavirus main protease. *J Med Chem* **49**, 5154-5161, doi:10.1021/jm060207o (2006).

3 Turlington, M. *et al.* Discovery of N-(benzo[1,2,3]triazol-1-yl)-N-(benzyl)acetamido)phenyl) carboxamides as severe acute respiratory syndrome coronavirus (SARS-CoV) 3CLpro inhibitors: identification of ML300 and noncovalent nanomolar inhibitors with an induced-fit binding. *Bioorg Med Chem Lett* **23**, 6172-6177, doi:10.1016/j.bmcl.2013.08.112 (2013).

4 de Wilde, A. H. *et al.* Screening of an FDA-approved compound library identifies four small-molecule inhibitors of Middle East respiratory syndrome coronavirus replication in cell culture. *Antimicrob Agents Chemother* **58**, 4875-4884, doi:10.1128/AAC.03011-14 (2014).
